# Metformin, empagliflozin and their combination modulate ex-vivo macrophage inflammatory gene expression

**DOI:** 10.1101/2022.06.20.496771

**Authors:** Adittya Arefin, Matthew C. Gage

**Affiliations:** Wolfson Institute for Biomedical Research, Division of Medicine, University College London, Gower Street, London, WC1E 6BT, United Kingdom; Department of Comparative Biomedical Sciences, Royal Veterinary College, 4 Royal College Street, London NW1 0TU, United Kingdom

**Keywords:** Macrophage, diabetes, inflammation, metformin, empagliflozin, combinations, anti-diabetes drugs

## Abstract

Type-2 Diabetes Mellitus is a complex, chronic illness characterized by persistent high blood glucose levels. Patients can be prescribed anti-diabetes drugs as single agents or in combination depending on the severity of their condition. Metformin and empagliflozin are two commonly prescribed anti-diabetes drugs which reduce hyperglycemia, however their direct effects on macrophage inflammatory responses alone or in combination are unreported. Here we show that metformin and empagliflozin elicit proinflammatory responses on mouse bone-marrow derived macrophages with single agent challenge, which are modulated when added in combination. In-silico docking experiments suggested that empagliflozin can interact with both TLR2 and DECTIN1 receptors and we observed that both empagliflozin and metformin increase expression of *Tlr2* and *Clec7a*. Thus, findings from this study suggest that metformin and empagliflozin as single agents or in combination can directly modulate inflammatory gene expression in macrophages and upregulate the expression of their receptors.

## Introduction

Type-2 Diabetes Mellitus (T2DM) is a complex, chronic illness characterized by persistent high blood glucose levels (Siu 2015). In 2017, 425 million people were reported to be suffering from T2DM, with this number projected to rise by 48% by the year 2045 to 629 million. The global yearly expenditure for healthcare costs of diabetes is projected to rise from 727 billon (2017) to 778 billion (2045) US dollar (International Diabetes Federation 2017).

Acute complications of T2DM include hypoglyceamia, diabetic ketoacidosis and hyperglycaemic hyperosmolar nonketotic coma (American Diabetes Association 2018, American Diabetes Association 2019). T2DM is strongly correlated with microvascular complications (including diabetic retinopathy, neuropathy and nephropathy) and macrovascular complications (such as cardiovascular diseases), which are the most common comorbidity associated with T2DM (Cade 2008). Intense management of blood glucose levels has been shown to reduce the microvascular complications associated with T2DM (UK Prospective Diabetes Study Group 1998, UK Prospective Diabetes Study Group 1998), but its impact on the outcome of cardiovascular diseases such as atherosclerosis is less clear (UK Prospective Diabetes Study Group 1998, Miller et al. 2013).

T2DM is a metabolic disease primarily characterized by decreasing sensitivity of cells in the body towards the endogenous insulin (insulin resistance) and decreasing insulin secretion (American Diabetes Association 2018) resulting in hyperglycaemia. Reduced insulin response may be due to a variety of factors including lipotoxicity, mitochondrial dysfunction, ER stress, hyperglycaemia and inflammation (Boucher et al. 2014).

### Macrophages play a significant role in T2DM progression

Macrophages are monocyte-derived phagocytic leukocytes of the innate immune system commonly associated with response to infection. However, macrophages also play a central role in the progression of T2DM through their ability to affect insulin response on metabolic tissues such as liver muscle and adipose through local inflammation (Liang et al. 2007).

Depending on the tissue microenvironment, monocytes can differentiate and polarize into proinflammatory (M1/classical) or anti-inflammatory (M2/alternative) macrophages or exist on a spectrum between these two extremes (Nagareddy et al. 2014, Woollard et al. 2010, Auffray et al. 2007, De Kleer et al. 2014, Yang et al. 2014). Recent studies have demonstrated that obesity and hyperglycemia promote myelopoiesis in mice and cause an expansion in the pool of circulating classical monocytes (Weisberg et al. 2006, Nagareddy et al. 2013). Classical short-lived monocytes produce inflammatory cytokines and these monocytes selectively penetrate the inflamed tissues (Nagareddy et al. 2014, Woollard et al. 2010, Auffray et al. 2007, De Kleer et al. 2014, Yang et al. 2014). This metabolic inflammation has become a major focus of research linking obesity, insulin resistance and T2DM (Esser et al. 2014), and is characterized by increased immune cell infiltration into tissues, inflammatory pathway activation in tissue parenchyma, and altered circulating cytokine profiles. TNFα, IL1β, IFNγ and IL6 are major inflammatory cytokines, which are upregulated in diabetes (Lachmandas et al. 2015), and atherosclerosis (Duewell et al. 2010) and are expressed in macrophages (Hu et al. 2021).

### Treating patients with T2DM

Management of T2DM is complex due to the chronic nature of the disease often progressing over decades combined with integrating the management and treatment of its associated comorbidities (Morris et al. 2014). Patients are advised to partake in lifestyle modifications including maintaining a healthy diet, regular physical activity and weight-loss (LeRoith 2019). Unfortunately, this is often ineffective (Morris et al. 2014) and so the patients are then prescribed different classes of anti-diabetes agents depending on their blood glucose levels and glycosylated hemoglobin level (% HbA1c) (American Diabetes Association 2019).

Common anti-diabetes drugs are aimed at reducing the hyperglycemia (International Diabetes Federation 2017, Maggi et al. 2019, American Diabetes Association 2018), by targeting tissues which directly impact blood glucose levels, for example metformin targets the liver by reducing hepatic glucose output (Maggi et al. 2019) and empagliflozin block glucose reabsorption from the kidneys (Maggi et al. 2019). The availability of different drugs to control hyperglycaemia provides opportunities for tailoring the treatment regimen according to the individual need of the patient. Typically, patients may be prescribed with a single drug or combination of drugs depending on the severity of their disease (American Diabetes Association 2019, American Diabetes Association 2018) in accordance with health research association guidelines such as the National Institute of Health Care Excellence (NICE) or American Diabetic Association (ADA). This approach imparts an increasing therapeutic burden on the patient - either in the form of dosage upregulation or additional medications (Chaplin et al. 2016, Espinoza et al. 2020).

The administration of long-term drugs is not without risks (Boye et al. 2020). These agents may reduce insulin resistance, increase insulin secretion and glucose absorption from blood (Chaudhury et al. 2017, Rakel et al. 2018). However, many of these agents may worsen the co-morbid metabolic disorders in T2DM patients (Maggi et al. 2019, Espinoza et al. 2020, Chaudhury et al. 2017, Rakel et al. 2018). For example, Thiazolidinediones are potent anti-hyperglycemic agents, yet have been associated with worsening of CVD and related mortality (Waller et al. 2018). Insulin secretagogues, for example sulfonylureas, meglitinides and DPP-4 inhibitors, have also been associated with higher CVD risk (Soccio 2014, Douros et al. 2018, Ou et al. 2016, Cosentino et al. 2018).

Recently the use of anti-inflammatory agents has shown improvement in hyperglycaemia control in T2DM patients and disease models (Esser et al. 2014, Kumar et al. 2015). Two common features of all of these agents are persistent reduction of inflammation (reduction in CRP levels in blood) and reservation of beta cell function, which collectively resulted in better hyperglycaemia management (Esser et al. 2014, Larsen et al. 2007, Larsen et al. 2009, van Asseldonk 2015, Vallejo et al. 2014, Cavelti-Weder et al. 2012, Hensen et. al 2013, Rissanen et al. 2012, Sloan-Lancaster 2013, Fleischman et al. 2007, Goldfine et al. 2008, Koska et al. 2008, Goldfine et al. 2010, Goldfine et al. 2013, Goldfine et al. 2013, Faghihimani et al. 2011, Bernstein et al. 2006). Thus, investigation of how immune cells such as macrophages respond to anti-diabetes agents requires closer attention. Further knowledge of any advantageous or disadvantageous effects of these drugs on the immune system, can be utilized to better treat the T2DM patients.

### Metformin and empagliflozin can affect macrophages responses

Several oral anti-diabetic agents have been reported to modulate macrophage polarization towards the M2 anti-inflammatory phenotype including metformin and empagliflozin (Stanley et al. 2011, Hattori et al. 2015, Hattori et al. 2018). However, the mechanisms underlying these effects are still poorly understood and may conflict. Metformin has been reported to promote M2 polarization (Xu et al. 2017) and antitumor or anti-angiogenic M1 polarization (Bastard et al. 2000). It has previously been shown in murine bone marrow derived macrophages (BMDM), that lipopolysaccharide (LPS) stimulated phosphorylation of p65 and JNK1 was decreased by metformin leading to reduced pro-inflammatory cytokine levels (Wang et al. 2018). In LPS-stimulated macrophages, reduction of ApoE expression has been reported to have been reversed by metformin via retarding nuclear translocation of NF-κB (Woo et al. 2014). It has also been reported that metformin can inhibit IL1β-stimulated release of IL6 and IL8 from macrophages, human smooth muscle cells and endothelial cells in a dose-dependent manner (Stavri et al. 2015, Isoda et al. 2006).

It has been recently suggested that the cardio-protective activity of empagliflozin (Pham et al. 2017) maybe due to its anti-inflammatory effect (Hattori et al. 2018). For example, empagliflozin has been reported to reduce the levels of C reactive protein and polarize macrophages towards the M2 phenotype in patients (Hattori et al. 2018). Empagliflozin reduces obesity-induced inflammation via polarizing M2 macrophages in white adipose tissue and liver (Pham et al. 2017) and empagliflozin has been reported to decrease M1 macrophages and increase in M2 macrophages in the liver and epididymal white adipose tissue of mice (Xu et al. 2017). In ex vivo experiments with macrophages stimulated with ATP, it has been observed that empagliflozin can attenuate NLRP3 activation (Xu et al. 2019).

It has been speculated that combining metformin with other drugs with anti-inflammatory effects on the macrophages (e.g., empagliflozin) may help to strengthen the therapeutic potential of metformin (Kim et al. 2020). While this combination remained to be investigated, it has been previously reported that drug combinations can enhance the anti-inflammatory and anti-oxidant activities in stimulated macrophages (Feng et al. 2021) and the combination of empagliflozin and gemigliptin has been seen to exert anti-inflammatory activity on LPS-stimulated macrophages (Vuong et. al. 2019). In this investigation we sought to define the direct immunomodulatory properties of metformin and empagliflozin on macrophages as single agents or in combination; reflecting a clinical approach to patient treatment.

## Methods and Materials

### Animal work and cell culture

All animal procedures and experimentation were approved by the UK’s Home Office under the Animals (Scientific Procedures) Act 1986, PPL 1390 (70/7354). BMDM were prepared from Ldlr-knockout mice and cultured as described before (Gage et al. 2019, Pourcet et al. 2016). In brief: L929 Conditioned Medium (LCM) was used as a source of M-CSF for the differentiation of the macrophages. After 6 days of differentiation, LCM-containing medium was removed, cells were washed three times in warm PBS and incubated in DMEM containing low endotoxin (≤10 EU/mL) 1% FBS and 20 μg/mL gentamycin without any LCM before being treated with anti-diabetes drugs (metformin; Sigma-Aldrich, empagliflozin; Generon) for concentrations and durations indicated.

### Gene expression analysis

Total RNA from was extracted with TRIzol Reagent (Invitrogen). Sample concentration and purity was determined using a NanoDrop™ 1000 Spectrophotometer and cDNA was synthesized using the qScript cDNA Synthesis Kit (Quanta). Specific genes were amplified and quantified by quantitative Real Time-PCR, using the PerfeCTa SYBR Green FastMix (Quanta) on an MX3000p system (Agilent). Primer sequences are shown in supplementary table S1. The relative amount of mRNAs was calculated using the comparative Ct method and normalized to the expression of cyclophylin.

### In silico molecular docking simulation

A high resolution (2.4 Å) 3D crystal structure of TLR2 (PDB ID: 3A7C) was selected from the protein data bank (Berman et al. 2003) and converted to PDB format. This structure was then processed to present the proper size, orientation, and rotations of the protein (Arefin et al. 2021). The processing was carried out in UCSF Chimera (version 1.14) (https://www.cgl.ucsf.edu/chimera/) to remove non-standard amino acids, water molecules, ligands and ions, add missing hydrogen atoms, and to perform energy minimization of the protein structure (Yang et al. 2012). The 3D structures of Zymosan (PubChem CID: 64689) and Empagliflozin (PubChem CID: 11949646) were obtained in sdf format from PubChem (Kim et al. 2021). Energy minimization of these compounds were performed in UCSF Chimera (version 1.14) based on the Gasteiger method to reduce the accumulative charge on ligands to zero (Gasteiger et al. 1978). After processing these molecules were saved as ‘mol2’ files for molecular docking. The docking experiments were conducted with the processed protein and ligands using PyRx 0.8 docking software (Dallakyan et al. 2014). The same process was repeated with a high resolution (2.8 Å) 3D crystal structure of DECTIN1 (PDB ID: 2CL8) to assess probable interaction with Zymosan (PubChem CID: 64689) and Empagliflozin (PubChem CID: 11949646).

### Statistical analysis

Results are expressed as mean (± SEM). Comparisons within groups were made using one-way ANOVA with Dunnett’s correction applied. P≤0.05 was considered statistically significant.

## Results

### Metformin promotes *Tnfa* and *Il1b* inflammatory gene expression in macrophages

In order to explore the direct effects of metformin on inflammatory gene expression in macrophages we examined mRNA expression of four well-established inflammatory genes (*Tnfa, Il1b, Il6* and *Ifng*) in mouse BMDM at physiologically relevant concentrations of 1 μM and 10 μM (Pernicova et al. 2014, LaMoia et al. 2020) at 2 hour and 24-hour timepoints. We observed that metformin increased mRNA expression of *Tnfa* after 2 hours at 1 μM (Fig. 1A, 1.41-fold, p= 0.002) and 10 μM (Fig. 1A, 1.36-fold, p= 0.002) and *Il1b* after 24 hours (Fig. 1F, 6.2-fold, p= 0.031).

**Figure 1:**
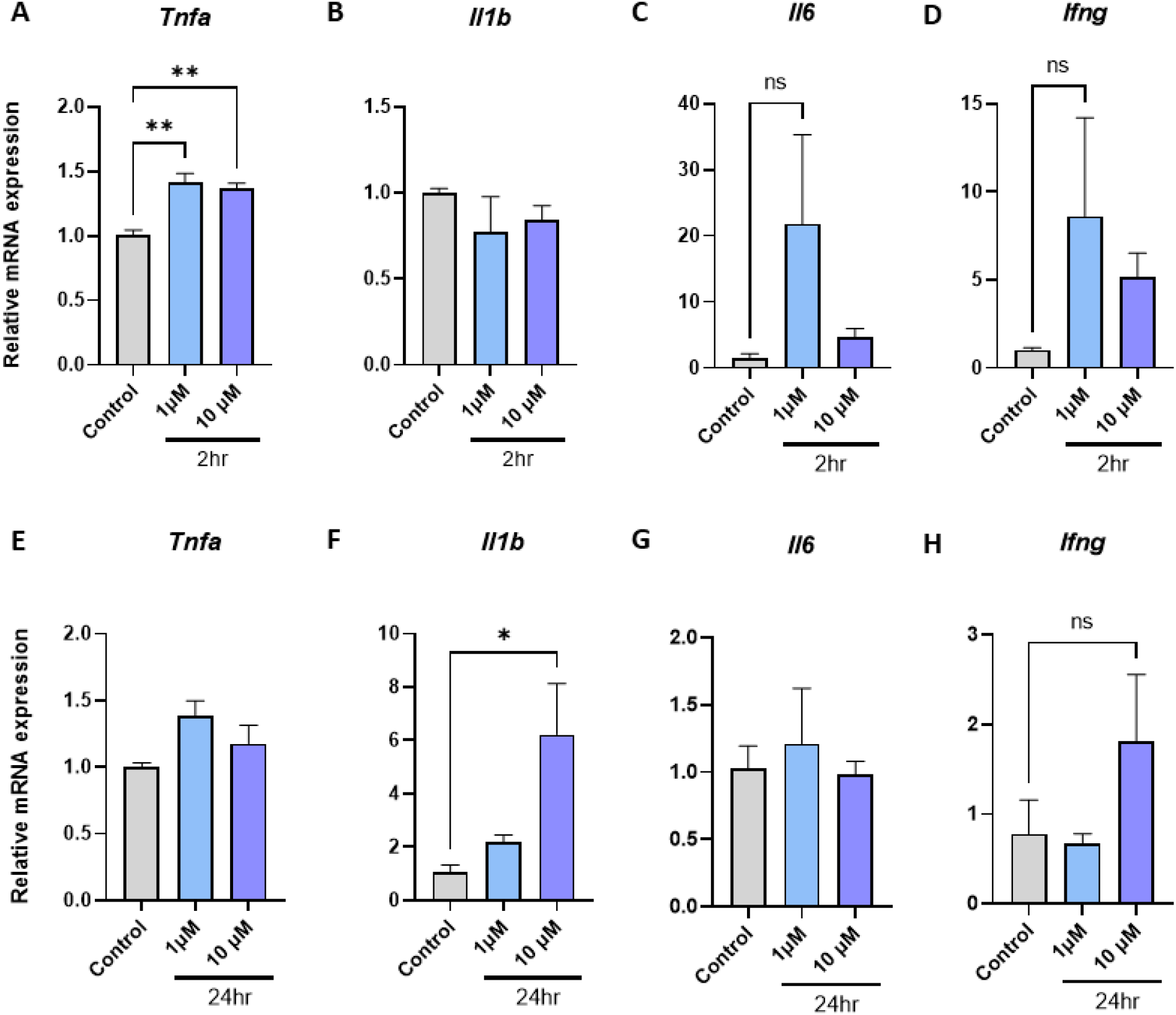
Metformin elicits direct proinflammatory gene expression in BMDM in a time and dose dependent manner. **(AD)** Metformin 2hr, **(E-F)** Metformin 24hr (n=3-4).

### Empagliflozin promotes *Tnfa, Il1b, Il6* and *Ifng* inflammatory gene expression in macrophages

To explore the direct effects of empagliflozin on inflammatory gene expression in macrophages we examined mRNA expression of the same four inflammatory genes at identical physiologically relevant concentrations (Boehringer Ingelheim International GmbH 2019) and timepoints. We observed that empagliflozin increased mRNA expression of *Tnfa* after 2 hours at 1 μM (Fig. 2A, 1.7-fold, p= 0.031), *Il6 at 1 μM (Fig. 2C, 13.7-fold, p= 0.037) and Ifng at 10 μM (Fig. 2D, 4.5-fold, p= 0.011) and Il1b* at 10 μM after 24 hours (Fig. 2F, 5.8-fold, p= 0.016).

**Figure 2:**
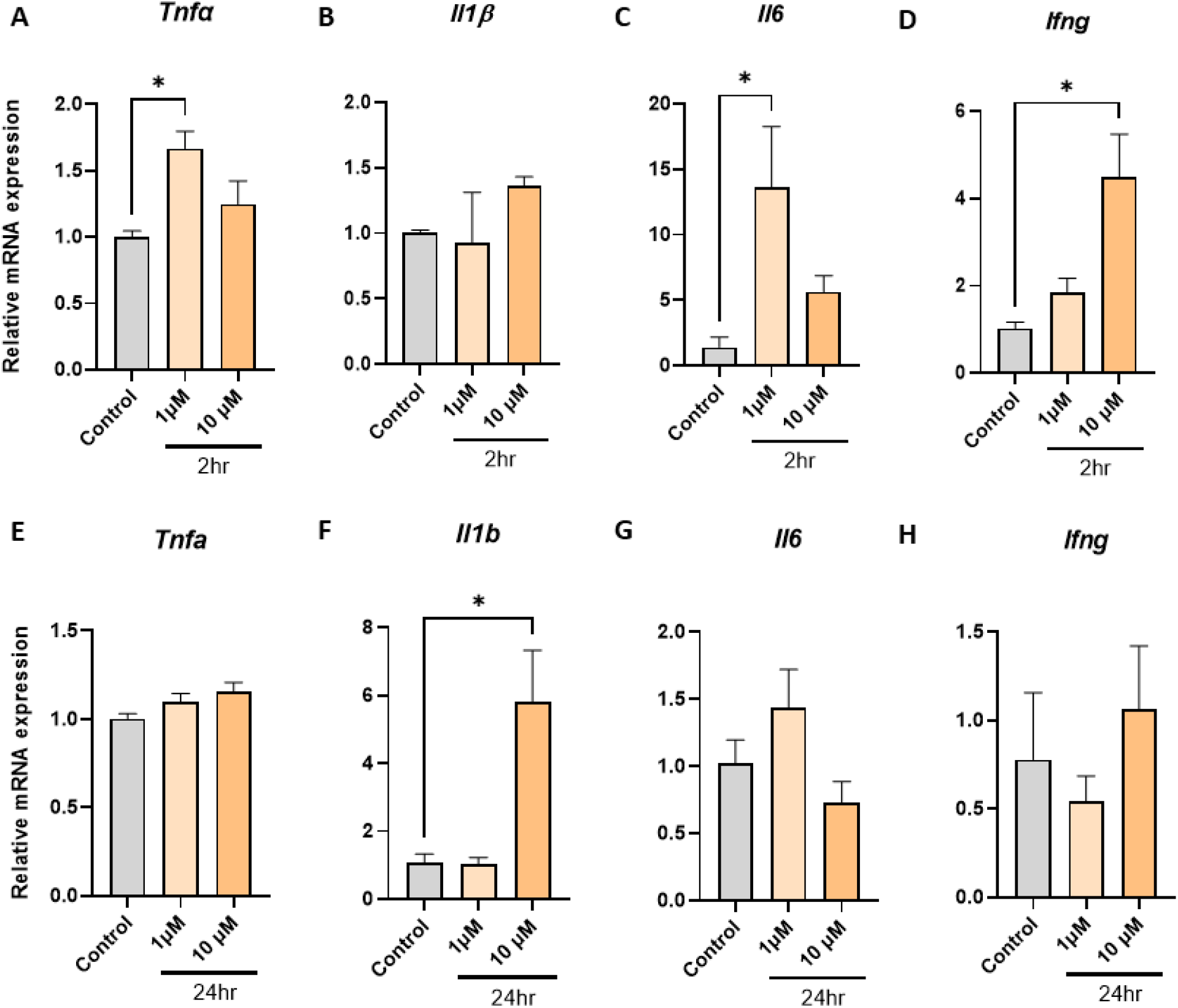
Empagliflozin elicits direct proinflammatory gene expression in BMDM in a time and dose dependent manner. **(A-D)** Empagliflozin 2hr, **(E-F)** Empagliflozin 24hr (n=3-4).

### Metformin and empagliflozin in combination have contrasting effects on macrophage inflammatory gene expression

As metformin and empagliflozin are commonly prescribed in combination, we next investigated how the combination of these drug might compare to the responses observed in the BMDM when they were added as single agents. We observed that in contrast to single drug responses, the combination of metformin and empagliflozin on *Tnfa* at 2h at 10 μM has no effect on mRNA expression (Fig. 3A), however after 24hr incubation, the levels of *Tnfa* mRNA expression were significantly increased (Fig. 3E, 1.4-fold, p= 0.019). Metformin and empagliflozin combination cancelled out the dramatic increase of mRNA expression of *Il1b* after 24 hours (Fig. 3F) and *Il6* after 24 hours (Fig. 3G) when compared to single agent responses (Fig. 1 and 2).

**Figure 3:**
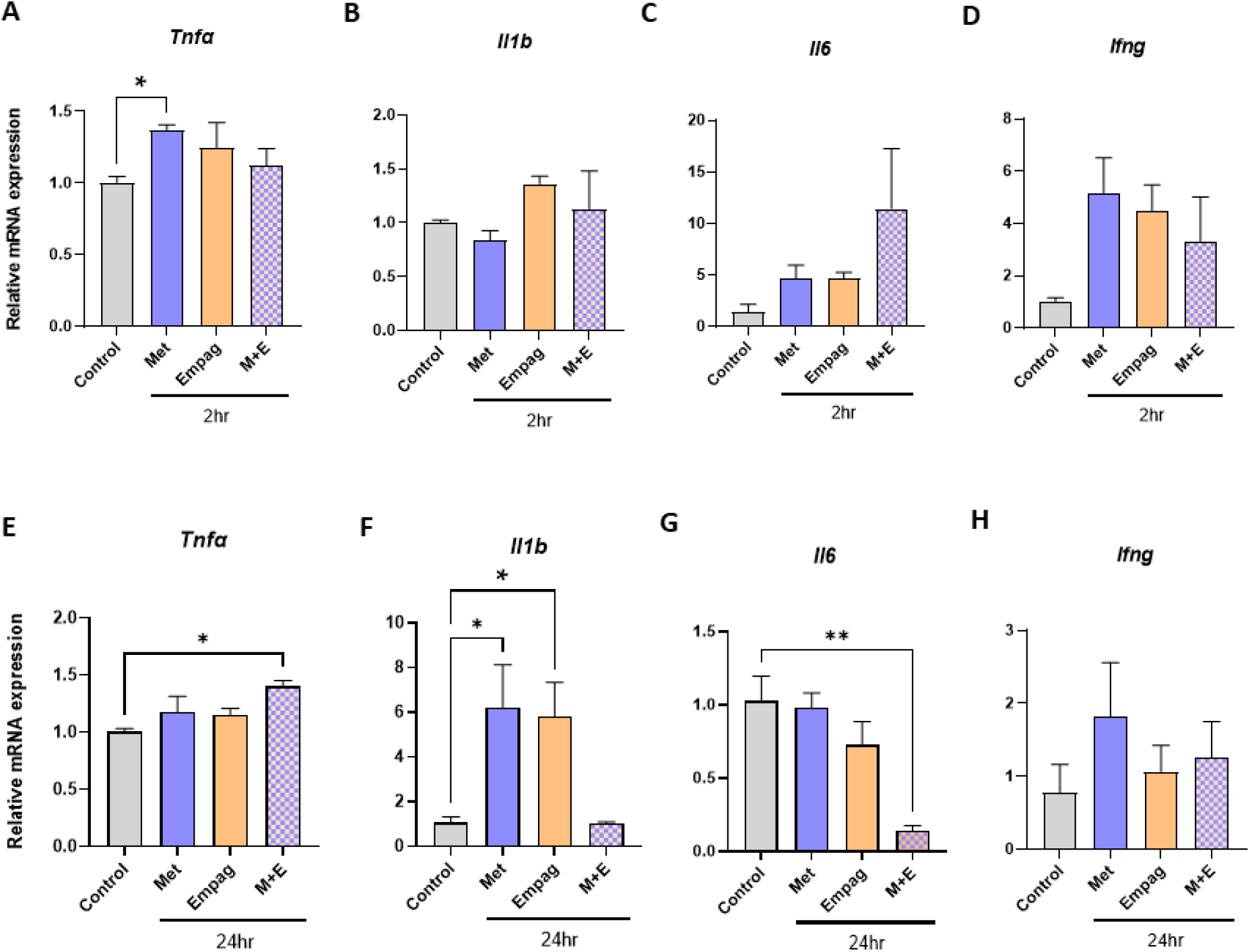
Metformin and empagliflozin in combination have contrasting effects on inflammatory gene expression in BMDM compared to single agents. **(A-D)** 2hr, 10 μM, **(E-H)** 24 hr, 10 μM (Met=Metformin, Empag = Empagliflozin, M+E = combination, n=3-4).

### Metformin and empagliflozin can interact with *Tlr2* and *Clec7a* and modulate their expression

The direct effects of metformin and empagliflozin on basal macrophage gene expression have not been reported previously. Inflammatory gene expression in macrophages can be induced through the macrophage’s expression of pathogen associated molecular pattern (PAMP) recognition receptors, which include the toll-like receptors (TLRs) (Oliveira-Nascimento et al. 2012) and DECTIN1 (InvivoGen 2019). Therefore, we speculated that the proinflammatory signalling we observed may be induced through these receptors. When investigating the structure of empagliflozin (PubChem CID: 11949646) we noticed that empagliflozin has a similar moiety to yeast zymosan (PubChem CID: 64689) (Fig. 4B). Zymosan is a well-established activator of inflammatory gene expression in macrophages through TLR2 and DECTIN1 (Li et al. 2021, Sato et al. 2003, Dillon et al. 2006)

**Figure 4:**
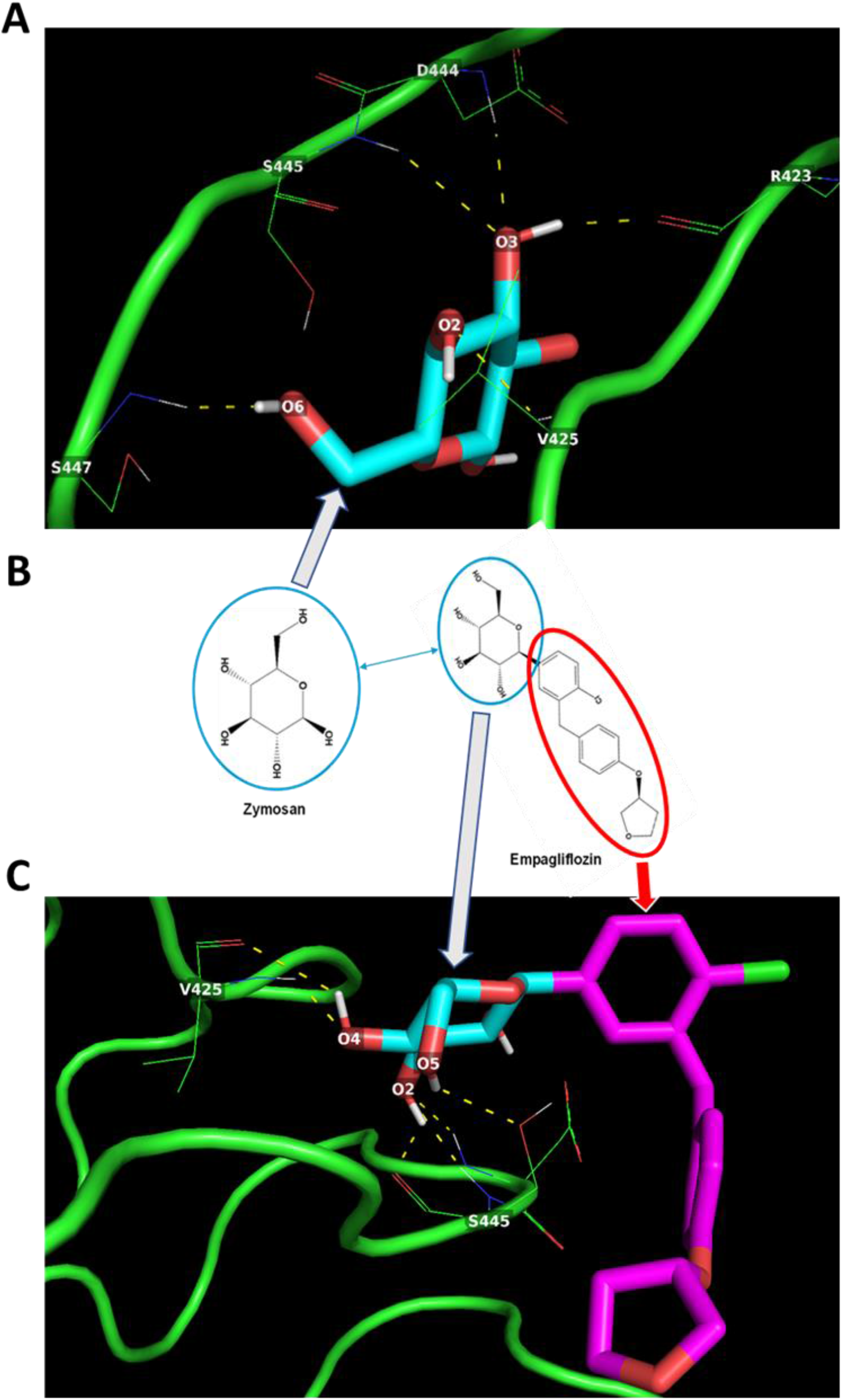
Potential zymosan and empagliflozin interactions with TLR2. (A) Zymosan may interact via multiple hydrogen bonds (dotted yellow lines) with R423, V425, D444, S445 and S447 amino acid residues of TLR-2. (B) Empagliflozin has a moiety identical to Zymosan. (C) the zymosan-like moiety may enable Empagliflozin to interact withV425 and S445 amino acid residues of TLR-2 via H-bond formation.

In-silico protein-ligand docking assessment suggests that both zymosan (Fig. 4A) and empagliflozin (Fig. 4C) could interact with the TLR2 through hydrogen bond interactions with amino acid residues R423, V425, D444, S445, S447 (Fig. 4). Remarkably, despite having multiple H-bond donor and acceptor groups, the H-bond formation between the residues of TLR2 and empagliflozin seemed to be facilitated only by the moiety identical to zymosan (Fig 4 and Table 1) with better predicted binding energy (−6.0 kcal/mol) than zymosan (−4.2 kcal/mol) (Table 1).

**Table 1:** Predicted protein-ligand interactions for TLR2-Zymosan and TLR2-Empagliflozin with binding energies from docking simulations.

| Target Protein | Ligand | Potential H-bond formation | Predicted Amino Acid residue interaction (number) | Predicted Binding Energy (kcal/mol) |
| --- | --- | --- | --- | --- |
| TLR2 | Zymosan | 5 | R423 (1)<br>V425 (1)<br>D444 (1)<br>S445 (1)<br>S447 (1) | - 4.2 |
| TLR2 | Empagliflozin | 6 | V425 (2)<br>S445 (4) | - 6.0 |

A similar result was observed during docking simulations with DECTIN1-Zymosan and DECTIN1-empagliflozin. Zymosan (Fig. 5A) can interact with DECTIN1 receptor through H-bond formation with H126, K128, S129, Y131, N159, E241 amino acid residues. On the other hand, empagliflozin can form H-bonds with Y131 and N159 amino acid residues of DECTIN1 (Fig. 5C). Again, the interaction of empagliflozin with DECTIN1 seemed to be facilitated by the moiety identical to zymosan (Fig 5 and Table 2) and yields better binding energy (−6.1 kcal/mol) than zymosan (−5.0 kcal/mol) (Table 2).

**Figure 5:**
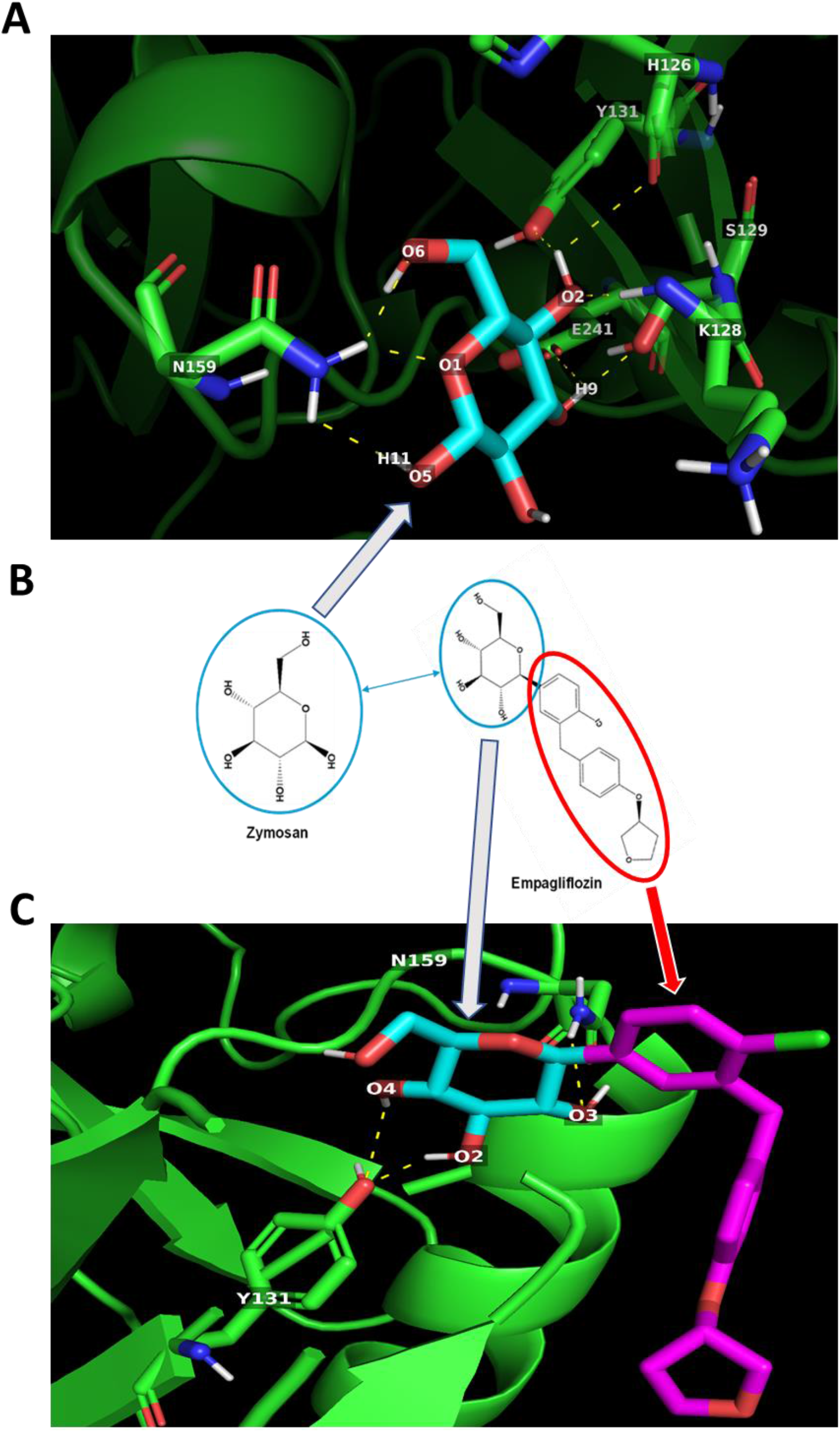
Potential Zymosan and empagliflozin interactions with Dectin-1. (A) Zymosan may interact via multiple hydrogen bonds (dotted yellow lines) with **H126, K128, S129, Y131, N159 and E241** amino acid residues of Dectin-1. **(B)** Empagliflozin has a moiety identical to Zymosan. (C) the Zymosan-like moiety may enable Empagliflozin to interact with **Y131** and **N159** amino acid residues of Dectin-1 via H-bond formation.

**Table 2:**
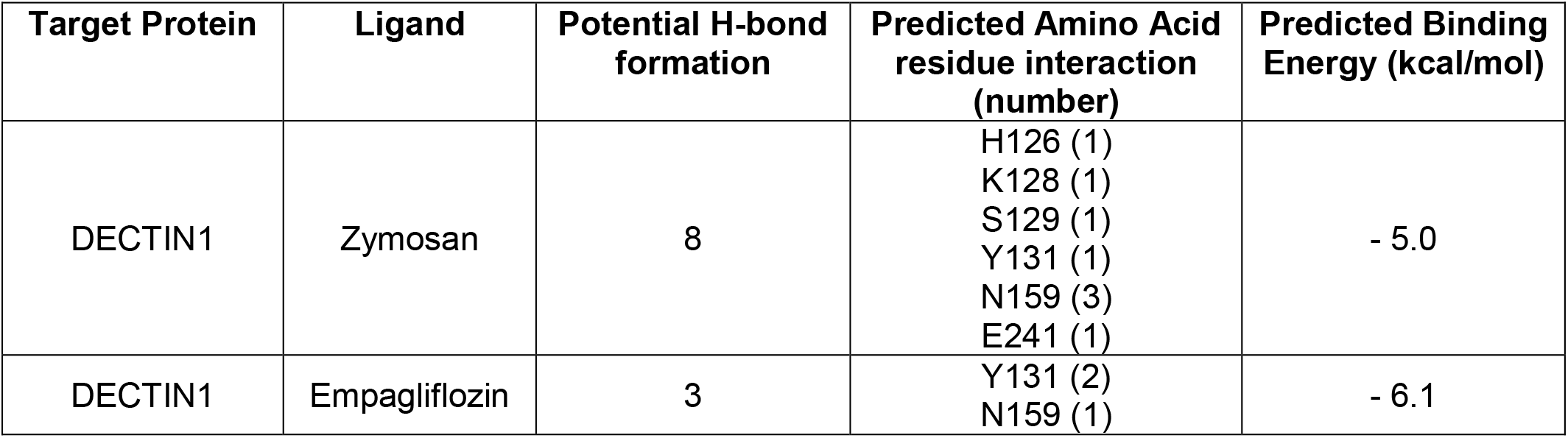
Predicted protein-ligand interactions for DECTIN1-Zymosan and DECTIN1-Empagliflozin with binding energies from docking simulations.

Follow-up experiments investigating the effects of metformin and empagliflozin either as single agents or in combination on *Tlr2* and DECTIN1 (gene symbol *Clec7a*) expression revealed that empagliflozin and metformin added as single agents at 10 μM increase *Tlr2* expression (Fig. 6A and C) at 2 hour (1.53-fold, p=0.0002; 1.38-fold, p=0.003) and 24-hour timepoints (1.37-fold, p=<0.0001; 1.26-fold, p=0.0005) respectively. Yet in combination, Tlr2 expression was less elevated (Fig 6A,1.24-fold, p=0.045) or negated (Fig.6C). Interestingly this mirrors the expression pattern of *Tnfa* after 2-hour exposure (Fig. 3A). On the other hand, exposures of 10 μM metformin, 10 μM empagliflozin and their combination increases *Clec7a* expression (Fig. 6B and D) at 24-hour (2.33-fold, p= 0.06; 2.23-fold, p= 0.08) respectively.

**Figure 6:**
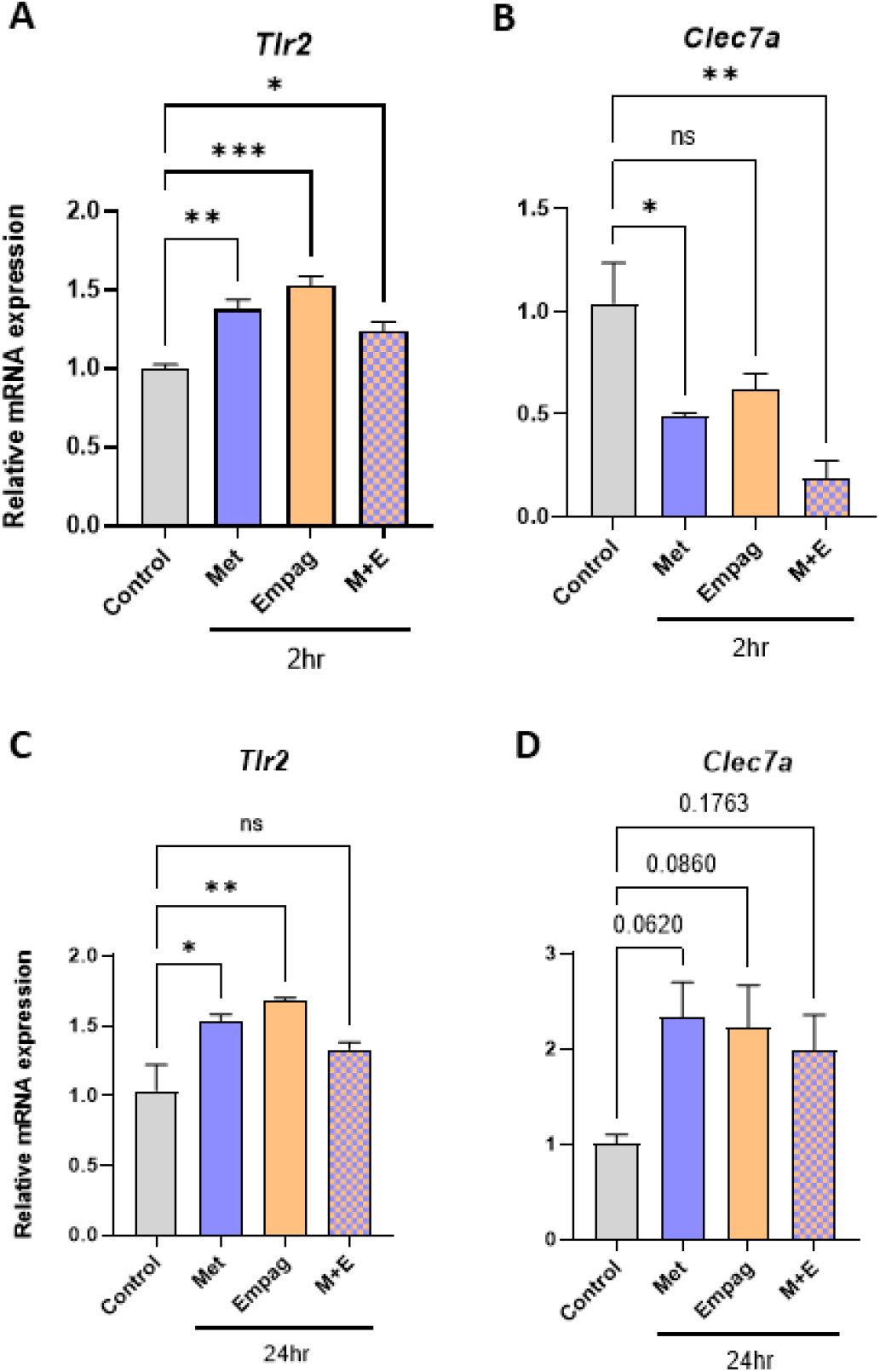
Metformin and empagliflozin in as single agents or in combination have contrasting effects on inflammatory gene expression in BMDM compared to single agents. **(A and B)** 2hr, 10 μM, **(C and D)** 24 hr, 10 μM (Met=Metformin, Empag = Empagliflozin, M+E = combination, n=3-4).

## Discussion

Depending on the severity of their disease, patients with type 2 diabetes may be treated with a monotherapy (such as metformin) or dual therapy combinations (such as metformin and empagliflozin combination) (Maggi et al. 2019). Macrophage-driven inflammation plays a significant role in the progression of type 2 diabetes (Lauterbach et al. 2017) and its associated comorbidities such as atherosclerosis (Lin et al. 2021). While reports are emerging of the indirect effect of anti-diabetes drugs on macrophages through polarization (Mathews et al. 2016) the direct responses of anti-diabetes drugs on these cells have remained unstudied. In this investigation we sought to determine the direct immunomodulatory properties of two of the most commonly prescribed anti-diabetes drugs metformin and empagliflozin on macrophages.

Metformin is a biguanide whose mode of action in reducing blood glucose is through reducing hepatic glucose production. Metformin does not require to be metabolised for its biological activity (Pernicova et al. 2014) and physiological plasma levels for its biological activity are reported to be between 1 μM to 40 μM with a half-life of 6.5 hours (LaMoia et al. 2020). Empagliflozin is a SGLT2 inhibitor whose mode of action is to block glucose reabsorption in the kidney. The physiological plasma levels for the biological activity of empagliflozin varies between 1.87μM to 4.74 μM based on the administered dose (10 mg and 25 mg respectively) and it is excreted from the body in an unchanged form after activity. The half-life of empagliflozin 12.4 hours (Boehringer Ingelheim International GmbH 2019). Therefore, to ensure the clinical relevance of our experiments we used metformin and empagliflozin at 1 μM and 10 μM for 2 hour and 24 hours to determine their direct immunomodulatory effect on murine bone marrow derive macrophages. Murine BMDM from LdlrKO mice are a well-established model for investigating macrophage responses in cardiometabolic settings (Gage et al. 2018, Neuhofer et al. 2014, Pendse et al. 2009, Dupasquier et al. 2007). Exposing BMDM to metformin at 1 μM and 10 μM for 2 hours increased the mRNA expression of *Tnfa* (Fig. 1A) and 24-hour exposure at 10 μM significantly increased the mRNA expression of *Il1b* (Fig. 1F). Exposing BMDM to empagliflozin also induced *Tnfa* expression at 1 μM within 2 hours (Fig. 2A), *Il1b* mRNA expression was significantly increased after 24 hours (Fig 2F). Significant increases in mRNA expression were also observed with *Il6* at 1 μM within 2 hours (Fig. 2C), and *Ifng* within 2 hours at 10 μM (Fig. 2D). Therefore, within the first 24 hours after physiologically relevant concentrations of metformin or empagliflozin exposure, several major inflammatory genes were observed to be up regulated.

Tnfa, Il1b and Il6 are activated through TLR signalling (Grassin-Delyle et al. 2020). Therefore, we speculated that the proinflammatory signalling we observed may be induced through these receptors. When investigating the structure of empagliflozin (PubChem CID: 11949646) we noticed that empagliflozin has a similar moiety to yeast zymosan (PubChem CID: 64689) (Fig. 4B). Zymosan is a well-established activator of inflammatory gene expression including *Tnfa* and *Il1b* in macrophages (Li et al. 2021, Sato et al. 2003, Dillon et al. 2006) through toll-like receptor 2 (TLR2) and DECTIN1 (mouse gene symbol *Clec7a*) we speculated the drug-receptor interaction may be TLR2 and DECTIN1 mediated. To test this hypothesis, in-silico molecular docking experiments were performed with crystal structures of TLR2 (Fig. 4) and DECTIN1 (Fig. 5) and the molecules zymosan and empagliflozin. The docking simulations not only suggested that empagliflozin can interact with both TLR2 and DECTIN1 receptors by similar amino acid residue interactions [Table 1 and 2] but also yielded better predicted binding energies for both the receptors compared to zymosan [Table-1 and 2]. These in silico docking experiments also revealed that only the zymosan-moiety in the empagliflozin chemical structure was predicted to be able to interact with TLR2 (Fig. 4B and C) and DECTIN1 (Fig. 5B and C) receptor amino acid residues. Collectively, these observations indicate towards a probable recognition of pathogen associated molecular pattern (PAMP) in the empagliflozin chemical structure by the macrophages. Ligand-receptor binding often modulates mRNA expression of the receptors involved (Papadopoulos et al. 2013). Further investigation revealed that empagliflozin modulates *Tlr2* and *Clec7a* mRNA expression (Fig. 6) in BMDM within the same timeframes observed for inflammatory gene expression lending support to their possible interaction.

Regarding the possible mechanism of metformin’s upregulation of the inflammatory genes observed; there is little published literature regarding metformin’s direct effect on macrophages. Metformin has historically been characterised by its ability to reduce hepatic glucose production through the transient inhibition of the mitochondrial respiratory chain complex I (Pernicova et al. 2014, LaMoia et al. 2020) and activation of the cellular metabolic sensor AMPK (Vasamsetti et al. 2014). Under physiological conditions metformin exists in a positively charged protonated form which may rely on different isoforms of the organic cation transporters (OCT) to enter the cell (Viollet et al. 2011, Higgins et al. 2012, Wu et al. 2018). However, over the last 15 years a much more complex picture of metformin’s roles is emerging, reflecting multiple modes of action which have AMPK independent mechanisms with the new findings varying depending on the dose and duration of metformin used (Rena et al. 2017). Our experiments revealed that metformin also upregulated *Tlr2* and *Clec7a* mRNA expression (Fig. 6) providing the opportunity for the mechanism behind this observation to the followed up in future investigations.

TNFα is an early response cytokine secreted by macrophages in response to pathogens which stimulates an acute phase immune response via pathogen associated molecular pattern (PAMP) receptors such as Toll like receptor 2 (TLR2) by regulating chemokine release and aiding further immune cell recruitment (Arango Duque et al. 2014). Our results suggest that macrophages upregulate *Tnfa* expression after being exposed to single antidiabetic agents (Fig 1A and 2A) A similar increase was also observed after 24-hour exposure (Fig.1E), however at this later timepoint the increase was not statistically significanct, possibly reflecting the more immediate nature of the TNFα response. The difference in effects observed at the higher concentration of 10 μM resembles typical responses observed through PAMP receptor stimulation whereby higher doses of PAMPs lead to a more intense immune response (Bauer et al. 2018. Makimura et al. 2006). Similar to TNFα, IL1β is also a pyrogenic cytokine produced by macrophages to initiate an inflammatory response to stimuli in its microenvironment. IL1β also regulates cytokine release acting as a chemoattractant for recruitment of immune cells to the site of inflammation (Arango Duque et al. 2014). One key difference between the two cytokines is that IL1β is synthesized as a leaderless precursor that must be cleaved by inflammasome-activated caspase-1 and then secreted as a mature isoform (Latz et al. 2010). Thus, compared to TNFα secretion and action, IL1β secretion and action become evident at a later time point. Our results demonstrate a similar pattern with exposure to single antidiabetic agents as significant increases in *Il1b* expression are observed at the later 24-hour timepoint (Fig. 1F and 2F). IL6 is a pleotropic cytokine with both inflammatory (Fernando et al. 2014) and anti-inflammatory (Yasukawa et al. 2003) effects and shared regulation pathways with TNFα and IL-1β production and secretion (Arango Duque et al. 2014, Nackiewicz et al. 2014). It has been previously observed in murine macrophages that TLR2 activation results in NF-κB activation which leads to up-regulation of Il6 expression (Hunt et al. 2018). Our results suggest that the increases we observe to *Il6* mRNA expression (Fig. 1C and 2C) may also be TLR2 mediated. IFNγ primes macrophages for enhanced microbial killing and inflammatory activation by TLRs (Su et al. 2015, Wu et al. 2014, Hu et al. 2008). In response to classic TLR stimulators (e.g., LPS) macrophages produce IFNγ (Schleicher et al. 2004, Fultz et al. 1993). Our results also suggest simultaneous upregulation of *Ifng* and post TLR-activation *Tnfa* expression (Fig. 1A, 1D, 2A and 2D). In addition, it has been reported that TLR2 stimulation in macrophages can retard the effects observed at 24-hour exposure to IFNγ (Benson et al. 2009). Observations from our study suggest that post TLR-activation *Tnfa* levels remained upregulated at 24-hour exposure to the drugs or combination (Fig. 1E, 2E and 3E) and *Tlr2* expression also remained significantly upregulated (Fig. 6C), although the previously observed upregulation in *Ifng* expression were lost at 24-hour exposure (Fig. 1H, 2H and 3H). Thus, it is possible that the drugs metformin and empagliflozin, alone or in combination, have mounted potent TLR2 mediated initial response augmented with upregulated *Ifng* expression.

Our results may seem to contrast to studies which report anti-inflammatory properties of metformin (Hattori et al. 2015, Woo et al. 2014, Stavri et al. 2015, Isoda et al. 2006, Feng et al. 2021) and empagliflozin (Hattori et al. 2018, Xu et al. 2017, Xu et al. 2019, Kim et al. 2020, Lee et al. 2021). However, these studies either report (1) indirect systemic anti-inflammatory effects which may be due to confounding factors such as reductions in hyperglycaemia (Hattori et al. 2015, Hattori et al. 2018, Woo et al. 2014, Isoda et al. 2006, Pham et al. 2017, Feng et al. 2021) or (2) polarizing effects (Xu et al. 2017, Wang et al. 2018, Pham et al. 2017, Xu et al. 2017, Vasamsetti et al. 2014). Our results show the direct effects of metformin and empagliflozin on macrophages.

As metformin and empagliflozin are often administrated in combination to patients with type 2 diabetes (American Diabetes Association 2018), we continued our investigation by exploring the effects of these drugs at 10 μM and at 2 hour and 24-hour time points. We observed that when added in combination the pro-inflammatory effects observed with single drug exposure at 2-hours were negated (Fig 3A and 3F). A similar pattern of differential modulation was seen with 24-hour exposure for *Tlr2* expression (Fig 6A). The mechanism of these reduced responses with metformin and empagliflozin combination may be due to these drugs being recognised by the same set of pattern recognition receptors and leading to competitive inhibition or development of tolerance due to sequential or simultaneous treatment with multiple or higher doses of PAMP (Bauer et al. 2018).

Surprisingly the exposure to combination of drugs significantly increased *Tnfa* mRNA expression at 24 hours (Fig. 3E) and the same combination significantly decreased *Il6* mRNA expression at 24 hours (Fig. 3G). Our data highlight the complexities of individual-gene macrophage inflammatory response regulation; we show a clearly coordinated proinflammatory response mediated by several genes to a single agent challenge (Figs 1 and 2) – which can be negated (Fig. 3A and 3F) or amplified (Fig. 3E) when challenged by a combination of those same agents (Fig. 7).

**Figure 7:**
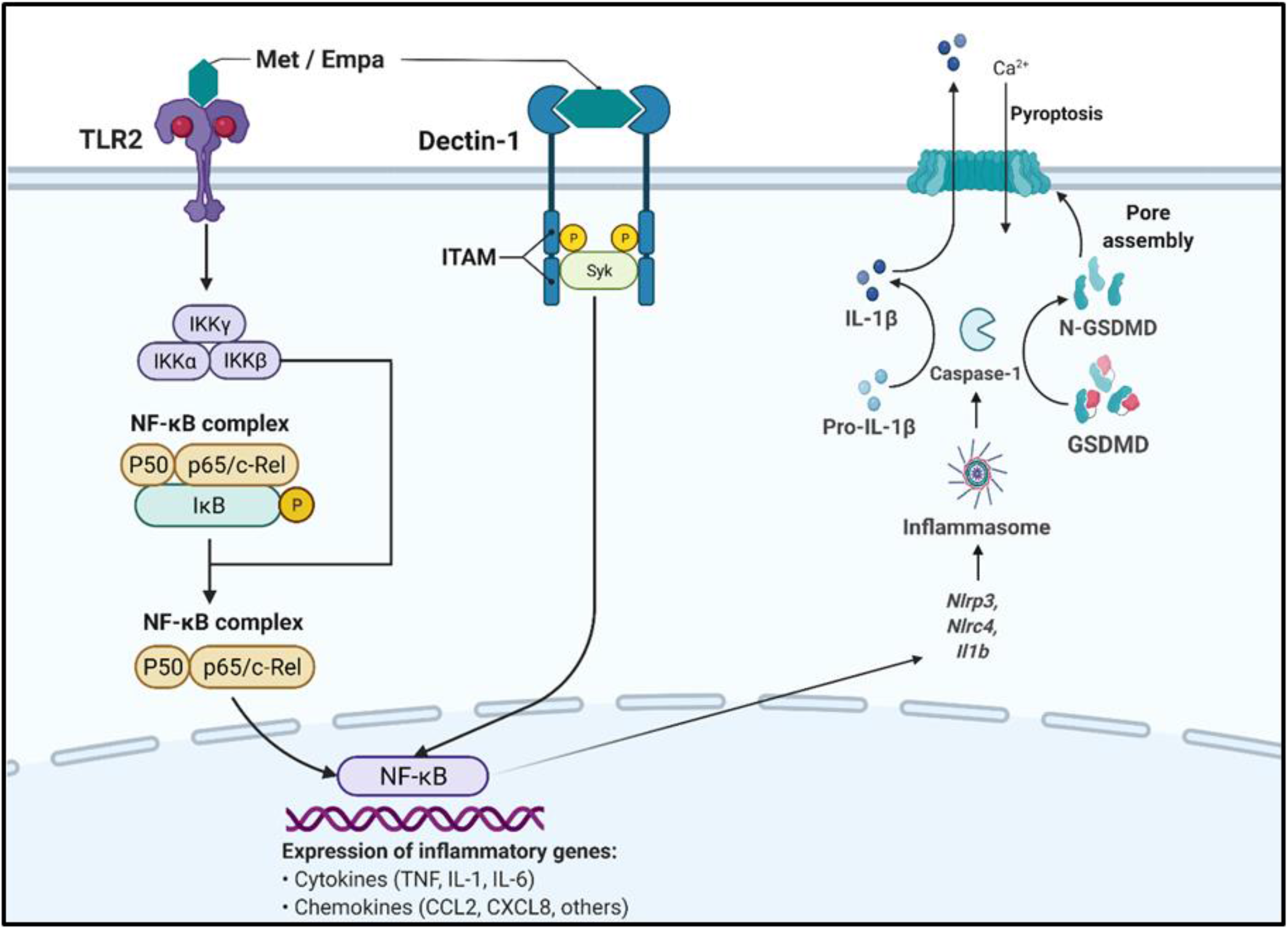
Schematic summarising the potential interaction of metformin and empagliflozin with TLR2 and Dectin-1 and how they may modulate macrophage inflammatory responses (met=metformin, Empa = empagliflozin). Created in BioRender.Com.

## Conclusion

In this investigation we sought to determine the direct immunomodulatory properties of two of the most commonly prescribed anti-diabetes drugs metformin and empagliflozin on macrophages. Murine bone marrow derived macrophages were exposed to clinically relevant concentrations and durations of metformin or empagliflozin in single doses and in combination. Our data show as single agents both metformin and empagliflozin elicit inflammatory responses in BMDM through cytokine and receptor expression and these responses are altered when the drugs are added in combination.

## Supplementary Materials

**Table S1:**
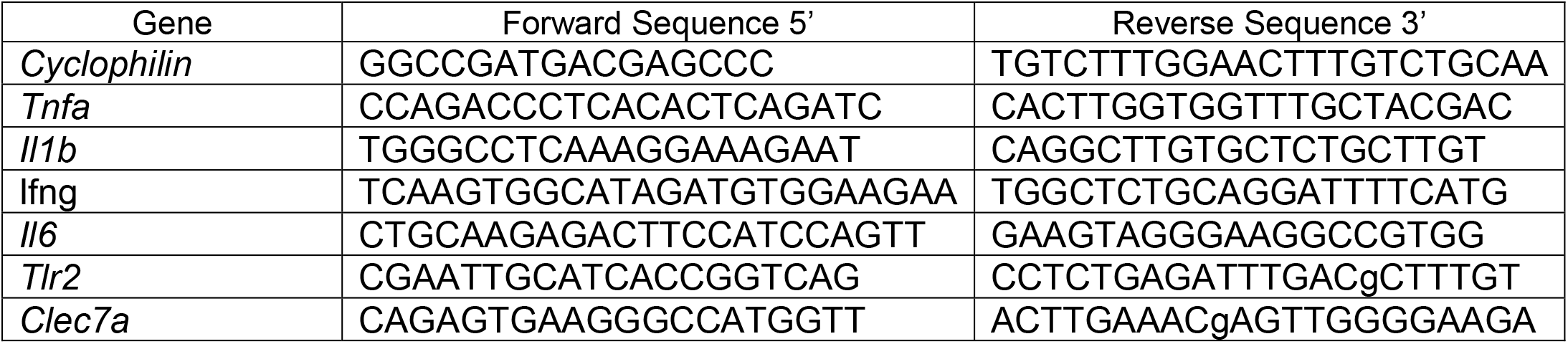
RT-qPCR Primers

## Acknowledgements

The authors would like to thank Inés Pineda-Torra for access to her laboratory and equipment and protocols and Gwladys Chabrier for her training and practical supervision of A.A.

## Author Contributions

Conceptualization A.A and M.C.G; Methodology A.A and M.C.G. Formal analyses A.A and M.C.G; Investigation A.A and M.C.G; Resources M.C.G, Data Curation A.A and M.C.G; Writing–Original Draft Preparation A.A and M.C.G; Writing-Review & Editing A.A and M.C.G; Visualisation A.A and M.C.G.; Supervision M.C.G; Project administration M.C.G. Funding acquisition M.C.G.

